# Translating genotype data of 44,000 biobank participants into clinical pharmacogenetic recommendations: challenges and solutions

**DOI:** 10.1101/356204

**Authors:** Sulev Reisberg, Kristi Krebs, Mart Kals, Reedik Mägi, Kristjan Metsalu, Volker M. Lauschke, Jaak Vilo, Lili Milani

**Affiliations:** Institute of Computer Science, University of Tartu, Tartu, Estonia; Software Technology and Applications Competence Centre, Tartu, Estonia; Quretec Ltd, Tartu, Estonia; Estonian Genome Centre, University of Tartu, Tartu, Estonia; Institute of Molecular and Cell Biology, University of Tartu, Tartu, Estonia; Department of Physiology and Pharmacology, Section of Pharmacogenetics, Karolinska Institutet, Stockholm, Sweden; Science for Life Laboratory, Department of Medical Sciences, Uppsala University, Uppsala, Sweden

**Keywords:** Pharmacogenetics, Pharmacogenomics, Biobank participants, Pre-emptive pharmacogenetic testing, whole genome sequencing, whole exome sequencing, HumanOmniExpress, Global Screening Array

## Abstract

**Purpose:** Biomedical databases combining electronic medical records, phenotypic and genomic data constitute a powerful resource for the personalization of treatment. To leverage the wealth of information provided, algorithms are required that systematically translate the contained information into treatment recommendations based on existing genotype-phenotype associations.

**Methods:** We developed and tested algorithms for translation of pre-existing genotype data of over 44,000 participants of the Estonian biobank into pharmacogenetic recommendations. We compared the results obtained by whole genome sequencing, whole exome sequencing and genotyping using microarrays, and evaluated the impact of pharmacogenetic reporting based on drug prescription statistics in the Nordic countries and Estonia.

**Results:** Our most striking result was that the performance of genotyping arrays is similar to that of whole genome sequencing, whereas exome sequencing is not suitable for pharmacogenetic predictions. Interestingly, 99.8% of all assessed individuals had a genotype associated with increased risks to at least one medication, and thereby the implementation of pharmacogenetic recommendations based on genotyping affects at least 50 daily drug doses per 1000 inhabitants.

**Conclusion:** We find that microarrays are a cost-effective solution for creating pre-emptive pharmacogenetic reports, and with slight modifications, existing databases can be applied for automated pharmacogenetic decision support for clinicians.

## INTRODUCTION

Genetic variation causing interindividual differences in drug response poses major problems for pharmacological therapy and drug development. In the recent decades a plethora of associations between genetic variants and treatment efficacy or adverse drug reactions has been identified^1^. However, the implementation of clinical pharmacogenomics is lagging far behind these discoveries^2^. Fast, accurate and cost-effective genotyping of genes involved in drug response is a crucial first step for the implementation of pharmacogenomics in clinical care. Ideally, the genotype data should already exist in an individual’s health record at the time when personalized treatment is necessary. The currently most widely used genotyping method is the array-based interrogation of (candidate) variants. However, due to recent progress in sequencing technologies, Next Generation Sequencing (NGS)-based methods, such as whole exome sequencing (WES) and whole genome sequencing (WGS) are becoming more prevalent. The advantage of the latter is that sequencing-based methods detect rare variants, which have been estimated to account for 30-40% of the functional variability in pharmacogenes^3^. Currently, multiple trials that evaluate the patient benefits of preemptive pharmacogenetic genotyping using the different methodologies are being conducted^4–6^.

For the translation of genetic testing results into treatment recommendations concerted efforts have led to the publication of genotype-based guidelines, for which strong evidence links genetic polymorphisms to variability in efficacy or risk for adverse reactions^7^. To account for the effect of allelic variation and haplotypes of genes relevant in drug response, the “star” (*) nomenclature system is most widely used^8^. For most genes covered by guidelines from the Clinical Pharmacogenomics Consortium (CPIC), comprehensive information tables have been prepared on how to define alleles on the basis of genetic variation, which facilitates the association of diplotypes with predicted phenotypes and thus their functional interpretation^8,9^. A collaborative effort is underway to develop a software tool (PharmCAT) for automated conversion of genotype information into CPIC guideline recommendations^10^.

Here, we provide an overview of the challenges and solutions for the translation of genotype and sequence data of 11 genes into pharmacogenetic diplotypes and recommendations for drug prescription. We leveraged genomic information of 44,448 Estonian Biobank participants genotyped by high density microarrays, WES or WGS and derived pharmacogenetic recommendations based on preexisting CPIC guidelines for 32 commonly prescribed medications. We find drastic differences in the predicted outcomes across genotyping platforms and demonstrate that WGS currently does not provide substantial additional actionable information regarding common pharmacogenetic alleles compared to the latest genotyping arrays. Importantly, these recommendations can be returned to biobank participants, or incorporated into their health records for the personalization of future treatment decisions. Furthermore, our analyses provide guidance for the optimal choice of clinical genotyping strategies.

## MATERIALS AND METHODS

### Overview of genetic data

The Estonian Biobank is a research-oriented biobank containing longitudinal data and biological samples, including DNA, for 5% of the adult population of Estonia. Participants of the biobank have signed a broad informed consent which allows the Estonian Genome Center to continuously update their records through periodical linking to central electronic health record databases and local hospital information systems^11^. Of the biobank participants, 8,132 have been genotyped using the HumanOmniExpress beadchip (OMNI) and 33,157 using the Global Screening Array (GSA) from Illumina. Furthermore, WES and WGS data is available for 2,445 and 2,420 participants, respectively (Figure 1A). Only 1,661 samples (3.7%) had been genotyped on more than one platform.

**Figure 1.**
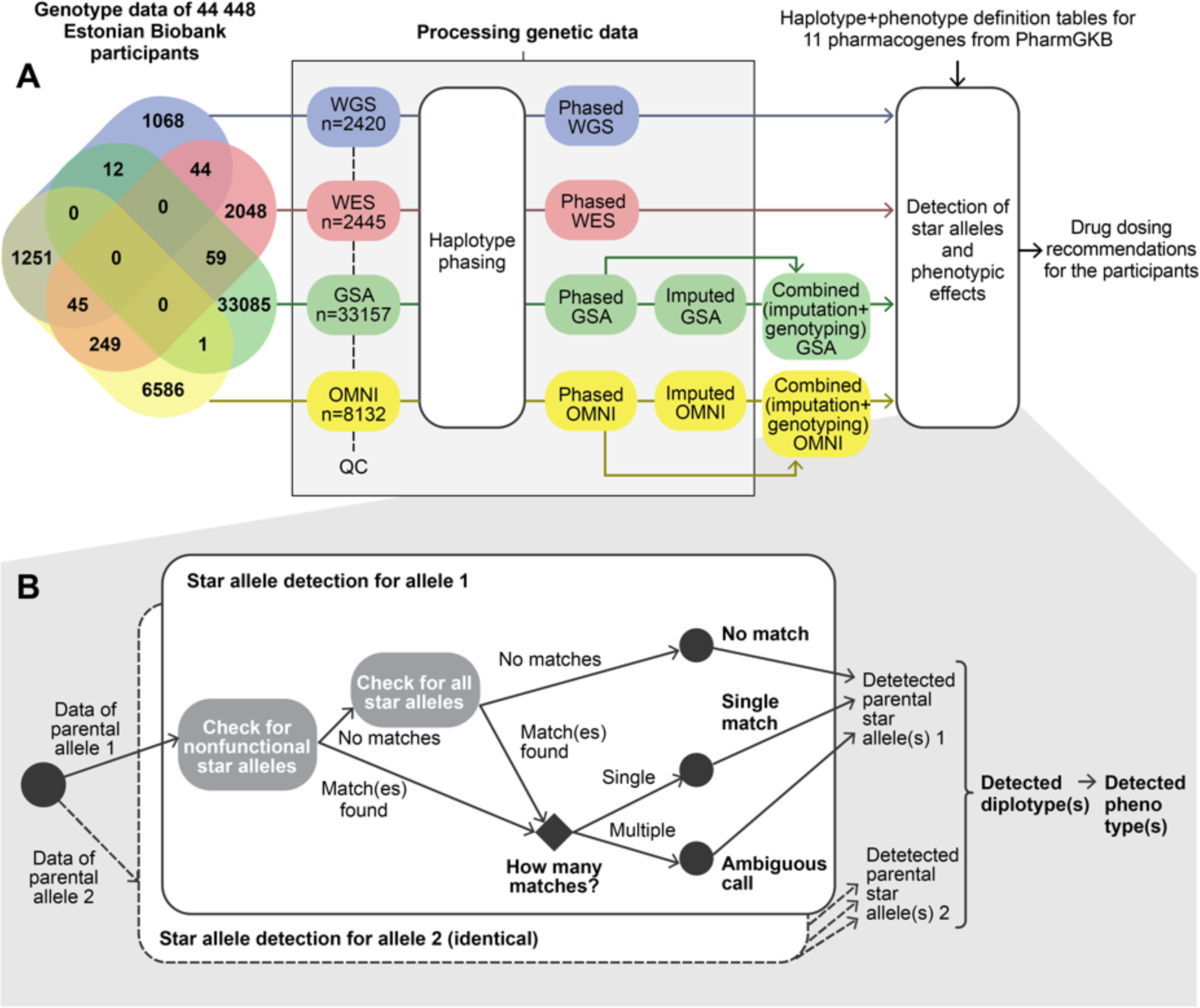
Pipeline for extracting pharmacogenetically relevant alleles from existing genotyping data. Panel A illustrates the different datasets, their overlap (Venn diagram) and how the data was processed. Panel B zooms into the detection of star alleles according to specific definition tables.

For whole genome sequencing, DNA samples were prepared using the TruSeq PCR-free kit, and sequenced on the Illumina HiSeq X using 150 bp paired-end reads at a mean coverage of 30x. WES samples were prepared using the Agilent SureSelect Human All Exon V5+UTRs target capture Kit according to the manufacturer’s recommendations, and sequenced on the HiSeq2500 at a mean target coverage of 67x. Sequenced reads were aligned against the GRCh37/hg19 version of the human genome reference using BWA-MEM^12^ v0.7.7; PCR duplicates were marked using Picard (http://broadinstitute.github.io/picard) v1.136, and the Genome Analysis Toolkit (GATK)^13,14^ v3.4-46 applied for further processing of BAM files and genotype calling. All insertion-deletions (indels) in the Variant Call Format (VCF)^15^ were normalized and multiallelic sites split using bcftools (https://samtools.github.io/bcftools/bcftools.html).

Samples were filtered based on high contamination (>5%), high proportion of chimeric alignment (>5%), low coverage (<20 **x** for WGS and below mean – 3SD for WES), low call rate for SNVs (<95%), discordance with genome-wide array data (>5%) and mismatch between phenotypic and genotypic sex. The following genotypes were set to missing: genotype quality <20, read depth >200 for WGS and <8 for WES, allele balance <0.2 or >0.8 for heterozygous calls. The GATK’s Variant Quality Score Recalibration (VQSR) metric was used to filter variants with a truth sensitivity of 99.8% for SNVs and of 99.9% for indels. Furthermore, variants with inbreeding coefficient <-0.3, quality by depth <2 for SNVs and <3 for indels, call rate < 95%, or Hardy-Weinberg equilibrium (HWE) P-value <1 **x** 10^−6^ were excluded.

A population-specific imputation reference panel^16^ contains the same subset of WGS individuals, but slightly different quality control parameters were used: only unrelated individuals were considered (IBD proportion <0.1); we excluded variants with call rate <90%, HWE P-value <1 **x** 10^−9^, multi-allelic variants, and low-complexity regions^17^. Finally, WGS data of 2279 Estonians and 1856 Finns were merged, where quality control was performed independently for each study.

The genotype calling for the GSA and OMNI arrays was performed using Illumina’s GenomeStudio V2010.3 software. The genotype calls for rare variants on the GSA array were corrected using the zCall software (version May 8th, 2012). After variant calling, the data was filtered using PLINK (v.1.90)^18^ by sample (call rate >95%, no sex mismatches between phenotype and genotype data, heterozygosity < mean +-3 SE) and marker-wise (HWE p-value >1 **x** 10^−6^, call rate >95%, and for the GSA array additionally by Illumina GenomeStudio GenTrain score >0.6, Cluster Separation Score >0.4). Before the imputation, variants with MAF <1% and C/G or T/A polymorphisms as well as indels were removed, as these genotype calls do not allow precise phasing and imputation.

The genotype data obtained on both arrays were separately phased using Eagle2 (v. 2.3)^19,20^ and imputed using the BEAGLE (v. 4.1)^21^ software implementing a joint Estonian and Finnish reference panel described above. Imputed genotypes with probabilities lower than 90% were filtered out. To call pharmacogenetic star alleles based on the microarray data we used genotyped variants together with imputed variants. In cases where the variant was both directly genotyped and imputed, the original genotype call was preferred. As a result of processing the genetic data, genetic information of all samples was converted into a joint VCF, where variant positions are aligned against the GRCh37/hg19 human genome reference.

### Pruning of Allele Definition Tables

To detect star alleles, we initially set out to use entire gene-specific Allele Definition Tables that have been prepared by the curators of PharmGKB and the Clinical Pharmacogenetics Implementation Consortium (CPIC) (https://www.pharmgkb.org/page/pgxGeneRef). We focused on the 11 clinically important pharmacogenes *CYP2C19, CYP2C9, CYP2D6, CYP3A5, CYP4F2, DPYD, IFNL3, SLCO1B1, TPMT, UGT1A1* and *VKORC1.* CPIC gene-specific tables of allele definitions, functionality, phenotype and frequency (downloaded on 17 Sep 2017) were used to first detect the pair of particular alleles for each gene and sample, and then estimate the corresponding phenotype. Out of the 356 variants in the CPIC tables used for defining the star alleles of these genes, 356 (100%), 307 (86%), 101 (28%) and 31 (9%) could potentially be detected by the WGS, WES, GSA and OMNI platforms, correspondingly, if the datasets contained individuals carrying the variants. However, as the allele definition tables are extremely large in some cases and are not accompanied with clear rules (e.g. decision trees) for prioritizing variants, direct application of the existing tables would result in a high proportion of ambiguous calls or no matches. Therefore, we first pruned the allele definition tables manually based on scientific evidence for functional effects of the variants.

First, we removed star alleles with unknown function or with unnecessary proxies (mostly suballeles) from *CYP2C19* (*35), *CYP2D6* (68 alleles, mostly suballeles), *DPYD* (*9A and *9B combined into *9) and *SLCO1B1* (32 alleles with unknown function), see **Table S1** for details. For *CYP2C19*2*, which is defined by 2 variants that are in complete LD (r^2^=1.0), we found that a single variant *(rs4244285)* is sufficient for its detection. Finally, we disregarded *CYP2D6* star alleles requiring gene deletions (*5) or duplications (star alleles with suffix “xN”) in the genotype and whole exome datasets, because detection of copy numbers of CYP genes is limited on these platforms. These filtering steps resulted in 239 variants remaining in the Allele Definition Tables. The final number of candidate star alleles that remained for each gene and data source after filtering is summarized in **Table S1**.

As there is no specific Allele Definition Table for typing of *HLA* alleles, we could not use the same pipeline for this region. However, to provide an overview of the relevant functional variability of the *HLA* region in the studied population, we used the SNP2HLA tool^22^ in the major histocompatibility complex region for the detection of *HLA* variants among individuals with WGS data.

### Pipeline for star allele and phenotype detection and analysis

For all of the samples we detected their possible star alleles by checking each star allele given in the Allele Definition Tables one by one and testing for the presence of defining variants for each allele. As this could result in several matching alleles due to missing data at certain positions, we found it reasonable to allow nonfunctional alleles to override other alleles. Therefore, we first check for the presence of variants defining nonfunctional star alleles only, and if none of these match, we test the remaining star alleles. In ideal cases, only a single star allele matched (“single match”) (**Figure 1B**). In some complex cases, the detected variants correspond to several star alleles (“ambiguous call”). Again, we reasoned that if one of the matching alleles was defined as “decreased function”, we let this override “normal function” alleles. Cases where an individual carried a combination of variants that did not have any corresponding star allele in the reference table were defined as “no match”.

For detection of *CYP2D6* large deletions, large duplications, and multiallelic copy number variants (CNV) in WGS data we used the Genome STRiP CNV discovery pipeline (version 2.00.1611)^23^ for 2269 deeply sequenced whole genomes. We used estimated information of *CYP2D6* CNVs together with our developed pipeline for star allele detection to assign CYP2D6 star-allele diplotypes. For detected duplications, we assumed an allele of the following order to be duplicated: *2>*1>*4, based on previous duplication frequencies in Europeans^24^.

For each sample, all possible diplotypes were constructed based on detected star alleles. The subsequent phenotype calling was based on PharmGKB’s diplotype-to-phenotype mapping tables.

The described pipeline was written as a custom Python script. The calculation part of the haplotype and diplotype detection was run in the High Performance Computing Center of the University of Tartu. The results of the allele, effect and phenotype detection were analyzed in R^25^ version 3.2.3 using the following packages: dplyr^26^, reshape2^27^ and ggplot2^28^.

Finally, we compared the obtained phenotype predictions to previously reported allele and phenotype frequencies of Caucasians (Europeans + North Americans). For this comparison, each sample was used once – WGS data was preferred over WES, GSA and OMNI. As a result, 2,420, 2,356, 33,086 and 6,586 samples were used from WGS, WES, GSA and OMNI data correspondingly. The results of 1,661 samples that had been sequenced/genotyped by more than a single method are compared in **Note S1**. For the WGS data, we also validated the non-structural star alleles and diplotypes of *CYP2D6* using an external tool Astrolabe (previously called Constellation)^29^. Furthermore, we estimated the potential clinical impact of the variants based on drug consumption statistics in Estonia (Annual statistical reports of the state agency of medicines), Finland (The Social Insurance Institution of Finland), Sweden (The National Board of Health and Welfare of Sweden), Denmark (statistics on the total sales of medicines in Denmark) and Norway (Drug Consumption in Norway 2012-2016).

## RESULTS

### Comparison of allele calls across four different genotyping platforms

We compared the pharmacogenomic predictions for biobank participants genotyped with any of four different microarray or sequencing platforms (**Figure 1**). Using the existing datasets combined with genotype imputation and phasing, we identified 100, 64, 61 and 43 different variants using GSA, OMNI, WGS and WES, respectively. Note that the larger number of variants in the microarray data is driven by more samples having been genotyped than sequenced. We assessed the imputation accuracy to be extremely high (99.96% matching genotype calls), which is described in further detail in **Note S1**.

Overall, the proportion of calls with no matches is very low (<0.05%) in all datasets. Ambiguous calls were slightly more problematic, ranging from 1.60% to 1.77%, except for WGS where 4.78% of the calls were ambiguous. This was caused by the co-occurrence of two star alleles of *UGT1A1* - *\*28* (defined by *g. 175490 CAT>CATAT*, that is not covered by the microarrays and did not pass QC in WES) and *\*80* (defined by *rs887829*, detected in all datasets). While the variants are defined as separate star alleles in the Allele Definition Tables, WGS highlights the high LD between these two variants by detecting both of them in 34.4% cases for this gene, resulting in a new *\*28+80* allele. All remaining ambiguous calls are mainly found in *CYP4F2*, where in addition to *\*2* and *\*3* (both defined by single variants) both variants *\*2+3* are detected in 15.5% of the samples. Therefore, an ambiguous call does not necessarily mean a weakness of a genotyping method, but may indicate a novel allele instead.

For the remaining 98.0% of the samples, star alleles for each gene were unambiguously detected. **Figure 2 A-E** and **Figure 3 A-F** show the frequencies of the detected star alleles by genotyping method. The full table of the frequencies of the detected alleles, including ambiguous calls and no matches, is provided in **Table S2**.

**Figure 2.**
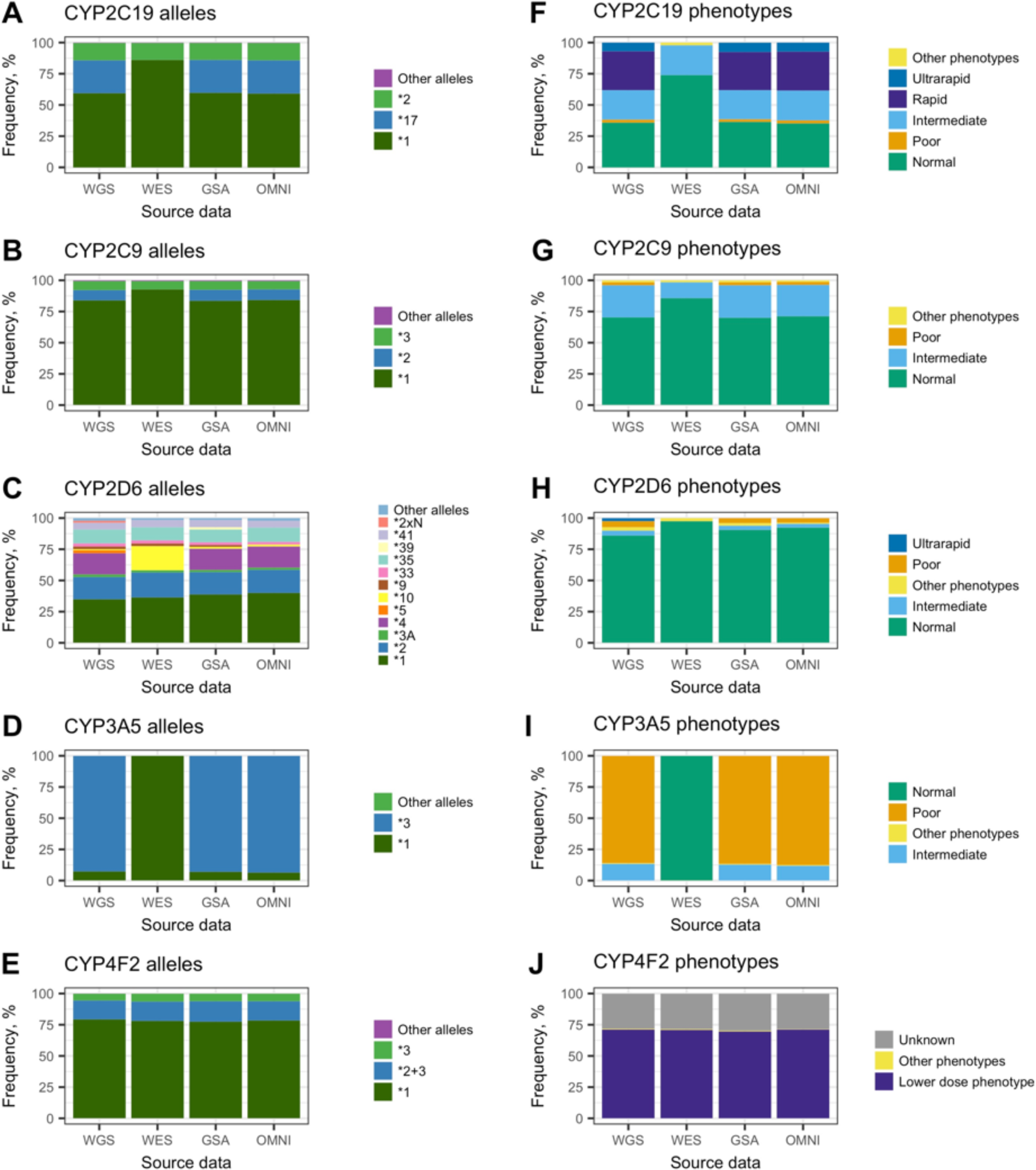
Frequencies of predicted alleles and phenotypes by CYP-gene and method. Alleles and phenotypes with frequencies below 2% are marked as “Other” for better visualization.

**Figure 3.**
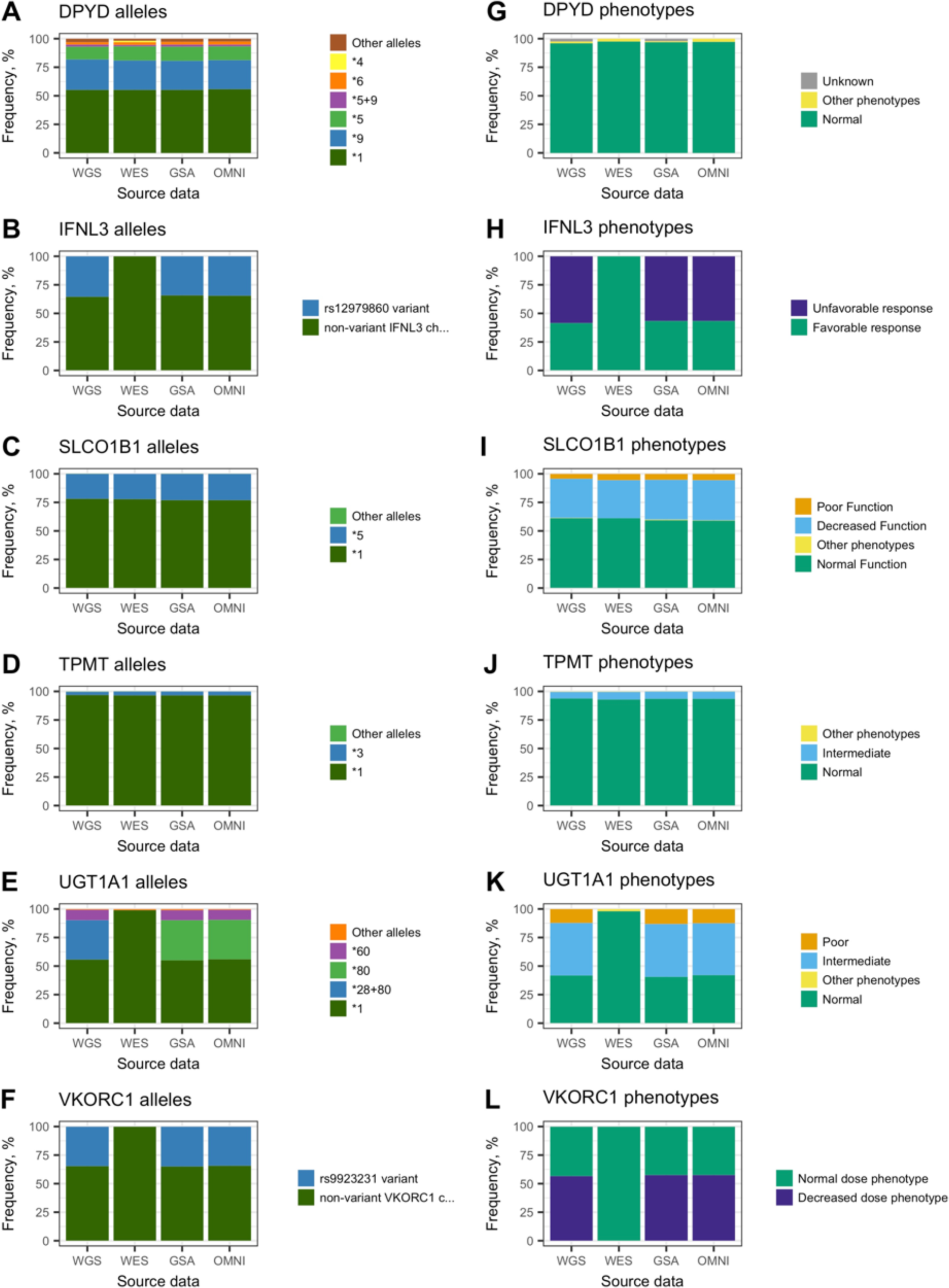
Frequencies of predicted alleles and phenotypes by gene and method for non-CYP genes. Alleles and phenotypes with frequencies below 2% are marked as “Other” for better visualization.

The figures clearly illustrate that the microarray based methods combined with imputation produce results that are very similar to WGS (except for *UGT1A1*, due to the reason described above). However, WES clearly underperforms. This is mostly caused by 11 star alleles that remain undetected due to variants falling outside the coding regions (see **Table S1** for details), but additionally, *CYP2C9*2* and *CYP2D6*4* could not be detected either, because the defining variants *rs1799853* and *rs3892097* did not pass QC.

To illustrate the proportion of rare variants detected in the 11 pharmacogenes under study, we assessed the frequencies of loss of function (LoF) and missense variants detected by WGS and WES in these genes (summary **Table S3**, full list given in **Table S4**). Altogether 89% (n=198) of the variants that we identified as putatively LoF or missense in the 11 pharmacogenes were rare with MAF <1% and 52% (n=102) of the variants were novel.

### Pharmacogenetic phenotype frequencies

Next, we used the called star alleles to derive actionable phenotypic predictions for all 11 analyzed genes (**Figure 2 F-J, Figure 3 G-L**). All diplotype frequencies are listed in **Table S5** and phenotype frequencies in **Table S6**. Similarly to the star allele calling the results are very similar for the different methods, with the exception of WES. From the perspective of implementing pharmacogenomics in the clinic, it is most crucial to accurately predict high risk phenotypes, i.e. individuals with other than normal drug metabolizing phenotypes and therefore require higher or lower dosing of a medication. The fraction of detected high-risk phenotypes for each gene and method are illustrated in **Figure 4**. Again, we observe that WES data is least suitable for pharmacogenomics, as a high proportion of high risk phenotypes remain undetected, except for *CYP4F2, DPYD, SLCO1B1* and *TPMT.* For *CYP3A5*, all phenotypes are detected as high-risk by WES, which is obviously incorrect. Therefore, we excluded WES results from the following analyses where we evaluated the presence of high risk phenotypes in 42,092 individuals, and found that non-standard dosing information is required based on at least one gene for 99.8% of the individuals.

**Figure 4.**
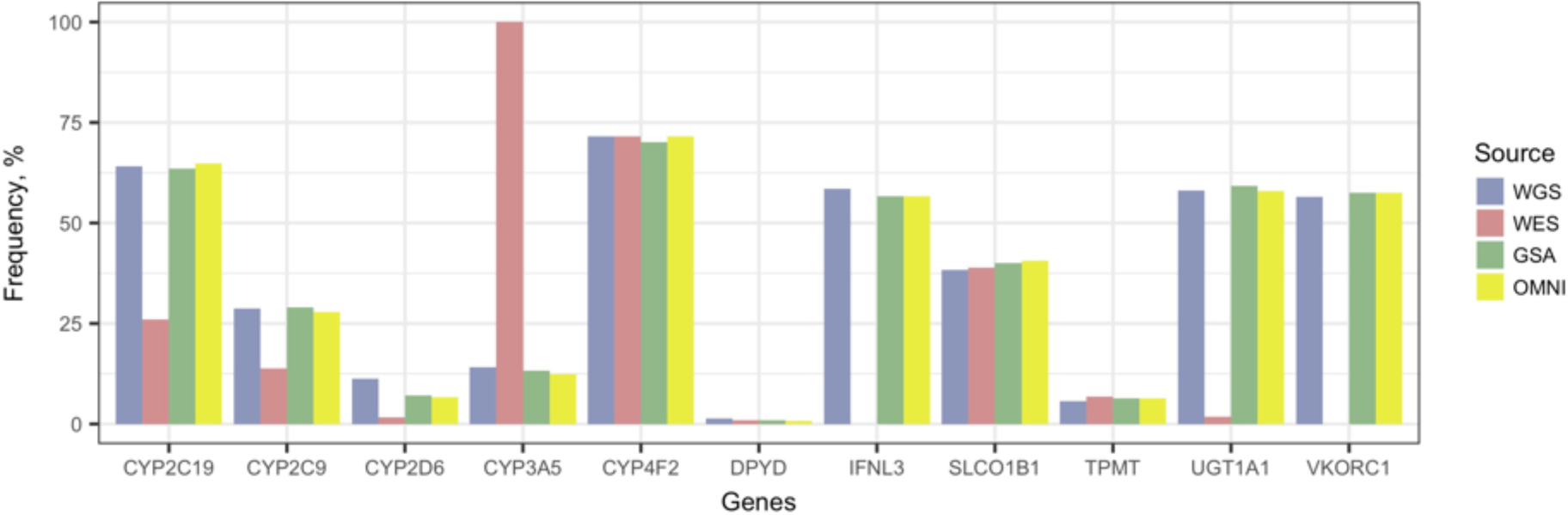
Fraction of high risk phenotypic predictions by gene and method. High risk phenotypes are defined as those that differ from normal and unknown phenotypes and would require a different drug dosing or recommendation.

The SNP2HLA tool allowed us to call 6-digit *HLA* haplotypes in the WGS dataset. Of the four high-risk phenotypes of the *HLA* region covered with CPIC guidelines^30–32^ we detected *HLA-B*57:01, HLA-B*58:01* and *HLA-A *31:01* alleles with carrier frequencies of 4.7%, 1.4% and 4.7%, respectively (**Table S3**). Since we were only able to call *HLA* alleles in the WGS data we could not compare the results between the different platforms.

We compared the results with frequencies reported in PharmGKB and by Muir *et al* ^33^ (see **Table S6** for details). In general, the frequencies of the detected alleles and phenotypes correspond to what has been reported previously. However, slight differences appear. For instance, there are significantly more *CYP2C19* rapid and ultrarapid metabolizers among Estonians (30.8% and 7.3%, respectively) compared to other Europeans (26.9% and 4.6%, respectively, p-values of one-proportion z-test 1.64×10^−72^ and 1.53×10^−155^). In addition, we compared the results we obtained for *CYP2D6* using our approach vs. a published tool Astrolabe. The same WGS data was used as input (2420 samples) for both methods, no structural variant information included. In 98% of the samples the detected alleles were identical, the discrepancies were mostly caused by *CYP2D6*59*, which is included in Astrolabe. We excluded this star allele from our candidate list due to sparse information about its suggested decreased function^24^. The overview of the comparison is illustrated in **Figure S1**.

### Relevance of detected phenotypes

Based on the dosing guidelines of CPIC, genetic variation in the 11 genes under study are associated with response to at least 32 currently prescribed medications (**Table S7**). *CYP2C19* affects the metabolism of drugs frequently used in the clinic^34^, and CPIC dosing guidelines are currently available for 10 active substances of these drugs. For this gene, we found that 2.2% of individuals in the studied cohort were poor metabolizers and 30.8 % and 7.3% rapid or ultrarapid metabolizers, respectively (**Table 1**). Thus, in total, 40.4% of the individuals in the Estonian population may be at risk for unwanted outcome or may need dosing adjustments when prescribed any of these 10 drugs. As shown in **Table 1**, the combined intake of medications associated with *CYP2C19* ranges from 17.62 - 66.83 DDD/1000 inhabitants per day in the Nordic countries and Estonia (Data from the Annual statistical reports, 2016).

**Table 1.** Frequencies of predicted high risk phenotypes within the studied cohort (WGS, GSA and OMNI data combined) and gene-related drug consumption statistics in European Nordic countries.

| Gene | Phenotype | % of individuals<br>(phenotype, source) |  |  | % of<br>individuals<br>(gene total) | Number of<br>drug active<br>substances<br>affected | DDD<sup>a</sup>/1000<br>inhabitants,<br>(min-max)<sup>b</sup> |
| --- | --- | --- | --- | --- | --- | --- | --- |
|  |  | WGS | GSA | OMNI |  |  |  |
| *CYP2C19* | Intermediate Metabolizer | 23.6 | 23.2 | 24.0 | 63.7 | 10 | 17.62 - 66.83 |
|  | Poor Metabolizer | 2.44 | 2.16 | 2.34 |  |  |  |
|  | Rapid Metabolizer | 31.2 | 30.7 | 31.2 |  |  |  |
|  | Ultrarapid Metabolizer | 6.86 | 7.40 | 7.23 |  |  |  |
| *CYP2C9* | Intermediate Metabolizer | 25.8 | 26.1 | 25.1 | 28.4 | 2 | 7.08 - 16.26 |
|  | Poor Metabolizer | 2.40 | 2.49 | 2.32 |  |  |  |
| *CYP2D6* | Intermediate Metabolizer | 3.93 | 3.23 | 2.96 | 7.32 | 16 | 9.16 - 15.92 |
|  | Poor Metabolizer | 4.96 | 3.93 | 3.67 |  |  |  |
|  | Ultrarapid Metabolizer | 2.36 | 0 | 0 |  |  |  |
| *CYP3A5* | Intermediate Metabolizer | 13.5 | 12.8 | 11.9 | 13.2 | 1 | 0 - 0.5 |
|  | Normal Metabolizer | 0.62 | 0.51 | 0.55 |  |  |  |
| *CYP4F2* | Higher dose phenotype | 0.29 | 0.36 | 0.33 | 70.5 | 1 | 7.02 - 16.04 |
|  | Increased CYP4F2 activity | 0.04 | 0.02 | 0.03 |  |  |  |
|  | Lower dose phenotype | 71.3 | 69.8 | 71.3 |  |  |  |
| *DPYD* | Intermediate Metabolizer | 1.36 | 0.90 | 0.87 | 0.92 | 3 | 0 |
|  | Poor Metabolizer | 0 | 0.006 | 0 |  |  |  |
| *IFNL3* | Unfavorable response | 58.5 | 56.7 | 56.7 | 56.8 | 3 | 0 - 0.23 |
| *SLCO1B1* | Decreased Function | 34.0 | 34.9 | 35.2 | 40.1 | 1 | 6.13 - 62.9 |
|  | Poor Function | 4.38 | 5.24 | 5.47 |  |  |  |
| *TPMT* | Intermediate Metabolizer | 5.54 | 6.37 | 6.33 | 6.40 | 3 | 0.32 - 1.41 |
|  | Poor Metabolizer | 0.21 | 0.07 | 0.08 |  |  |  |
| *UGT1A1* | Intermediate Metabolizer | 45.9 | 46.2 | 45.3 | 59.0 | 2 | 0 - 0.09 |
|  | Poor Metabolizer | 12.3 | 13.1 | 12.6 |  |  |  |
| *VKORC1* | Decreased dose phenotype | 56.5 | 57.5 | 57.5 | 57.4 | 1 | 7.02 - 16.04 |

<sup>a</sup> Drug daily dosage
<sup>b</sup> Min-max among Estonia, Finland, Sweden, Denmark, Norway
<sup>c</sup> Frequency calculated based on WGS only, as this was the only platform that detected *CYP2D6* ultrarapid metabolizers

Further, we also investigated the number of individuals with high risk variants that had been prescribed drugs associated with the specific genes. As seen in **Table S7**, as many as 12,254 individuals in the Estonian Biobank have actually had a prescription of at least one drug linked to *CYP2C19.* Out of these, 9,977 were analyzed in our study (WGS, GSA and OMNI) and 40.7% of them (n=4,059) are *CYP2C19* poor, rapid or ultrarapid metabolizers, and therefore may have needed dosing adjustments to improve treatment outcome. Based on the annual statistics of the Estonian Agency of Medicines, on average almost 5.5% (55 DDD/1000 inhabitants/day) of individuals in the population use at least one of the 32 drugs associated with the studied genes on a daily basis. For several Nordic countries, the numbers are even higher, the highest for Denmark with on average 15.8% of individuals in the population (158.2 DDD/1000 inhabitants/day) (**Table 1, Table S7**). Thus, existing data of biobank participants can be an untapped resource for improved and more cost-effective recommendations for drug treatment by translating existing genotype/phenotype data of pharmacogenes into guiding prescription recommendations. This illustrates the enormous innovative potential of biobanks in the whole process of the implementation of pharmacogenomics.

## DISCUSSION

In this study, we assessed the systematic detection of pharmacogenetic star alleles for Biobank participants genotyped on different microarray or sequencing platforms. As most of the pharmacogenes have star alleles defined by several variants that all need to be on the same parental allele, a crucial step in the process was genotype phasing prior to analysis. Although the PharmGKB tables for defining star alleles have been thoroughly curated, prefiltering of the Allele Definition Tables, as described in the Methods section, was essential for reasonable detection of star alleles. Many of the allele definitions include additional variants beyond the variant(s) causing the functional effects, which can compromise allele calling when searching for perfect matches. For example, in the original *SLCO1B1* star allele definition table, 20 alleles out of 37 require the occurrence of several mutations on the same allele, but in our dataset of 44,448 individuals, only a subset of these were actually detected on the same alleles, ruling out all possible star alleles and subsequently leading to “no matches” without prior filtering. The same applies for *CYP2D6*, where less than half of the alleles are currently of relevance^35^ and including too many unvalidated alleles would only result in unknown phenotypes. Challenges with these definition tables have been observed by others as well with an additional remark that the tables do not contain all of the alleles that are common in respective populations^24,36^.

We identified that 89% of the variants assessed in whole genome and exome sequencing data predicted to have functionally deleterious effects, are rare with MAF <1%. The proportion of rare variants detected in pharmacogenes has increased with the growing numbers of NGS studies^3,37-43^. Including rare variants with unknown function in pharmacogenetic reporting is objectionable, as their function and relevance are generally not well validated^2,9^ and care must be taken when including these in clinical implementation^44^. However, including rare variants in test panels and collecting data on these variants is still valuable for further research and development projects. In the absence of experimental characterization data, the functional impact of variants can be predicted using computational methods which are getting more and more precise with the increase in data that can be used for validation^45,46^.

With these prior filtering steps and by allowing non-functional star alleles to override other alleles, we were able to call star alleles of the genes under study for 98% of the individuals (Figure 2 and 3). The issue we faced with ambiguous calls (2%) due to the co-occurrence of two star alleles for *UGT1A1* and *CYP4F2* resulting in new “merged” star alleles, *\*28+80* and *\*2+3*, respectively, has been corrected for *UGT1A1* in the current updated version of PharmGKB (May 2018). The detection of several alleles on the same haplotype is not unexpected for such common variants, and combined with the challenges raised above, a functional variant driven approach might be more effective for calling of star alleles in general.

Our comparison of different genotyping and sequencing platforms highlights a known major shortcoming of WES for pharmacogenetic applications. Important alleles defined by variants in introns or promoters, such as *CYP2C19*17, CYP3A5*3* or *UGT1A1*28+80* are not interrogated by WES and thus lead to drastically different pharmacogenetic recommendations that affect 13 medications according to CPIC guidelines. Unlike microarray data, WES data cannot be subjected to classical imputation due to large gaps in the data. These problems could be overcome by combining WES with customized capture probes to provide a comprehensive cost-effective implementation of pharmacogenomics compared to WGS^47^. However, when the focus is exclusively on predefined alleles, genotyping arrays, which are currently at least 10 times cheaper than WES or WGS, are clearly a more cost-effective alternative that can generate results surprisingly similar to that of WGS. The versions of the arrays used in our study unfortunately do not allow the detection of *CYP2D6* copy number, which is the greatest but still limited drawback when compared to WGS (**Table 1**). As cost-effectiveness is still considered a major barrier for the clinical implementation of pharmacogenetics^48^, we suggest that current genotyping microarrays constitute the most cost-effective technology with acceptable accuracy. Several studies have found pre-emptive pharmacogenetic testing cost-efficient with per-patient savings ranging from 5,962-10,667 USD^49-51^, despite the reported costs of pharmacogenetic testing to be over 2,000 USD^49^. Thus, both genotyping and developing tools for translating pre-existing genome-wide genotype data into clinical recommendations can be considered very reasonable healthcare investments.

In conclusion, as the number of sequenced and genotyped participants in biobanks and clinical settings is growing rapidly in several countries, we now have a large amount of genetic information that could be translated into clinically actionable decisions tailoring medical therapy in the near future. By leveraging the existing genotype data of 44,448 individuals in the Estonian Biobank, we were able to determine that microarrays with imputed variants are a highly cost-effective tool for identifying thousands of individuals who need dosing adjustments for commonly prescribed drugs. In total, we found that as many as 99.8% of the individuals have a high risk phenotype requiring a non-standard dosing of a medication based on at least one gene, which is even larger than shown before^44,52^. Our approach of trying to define all possible star alleles in the majority of genes with CPIC guidelines allowed us to reveal the many challenges that arise in this process. The most crucial next steps we suggest are further revision of star allele definition tables based on existing haplotypes in different populations, an additional level of decision trees to prioritize variants causing nonfunctional alleles, and restricting the inclusion of rare alleles to functionally validated variants. We are confident that such developments built into automated decision support for clinicians will allow the implementation of pharmacogenomics at the point of care in a multidisciplinary manner^53^ and with greater impact.

## ACKNOWLEDGEMENTS

The study has been supported by the European Union through the European Regional Development Fund. The analysis of the data described in the article was carried out in the High Performance Computing Center of the University of Tartu. We would also like thank Meeli Mets for assistance with compiling the figures in the paper.

## Author Contributions

SR, LM and KK coordinated the study.

MK and RM prepared the genetic data and performed quality control.

SR, MK and RM phased the data and conducted genotype data imputation.

SR, KK, VL, LM, MK and RM designed the methodology.

SR wrote the software and ran the analysis.

SR, KK, LM and KM conducted the investigation.

SR prepared the figures.

SR, KK and LM wrote the manuscript.

SR, KK, LM, MK, RM, VL and JV reviewed and edited the manuscript.

## Supplementary Information

**Table S1. Size of star allele definition tables**

**Note S1. Variant calls were highly accurate**

**Table S2. Frequencies of the detected alleles**

**Table S3. The frequencies of predicted functional variants in 12 pharmacogenes (incl. HLA) identified in whole genome and exome sequencing data**. The table covers the frequencies of putative loss of function (LoF) and missense variants detected by sequencing. For *HLA* we evaluated the frequencies of well-known functional haplotypes included in CPIC dosing guidelines.

**Table S4. List of 198 putative loss of function (LoF) and missense variants detected in whole genome and exome sequencing data of 4,776 participants of the Estonian Biobank.**

**Table S5. Frequencies of the detected diplotypes**

**Table S6. Frequencies of the detected phenotypes**

**Table S7. Drug usage in Northern European countries**

---

**Figure S1.**
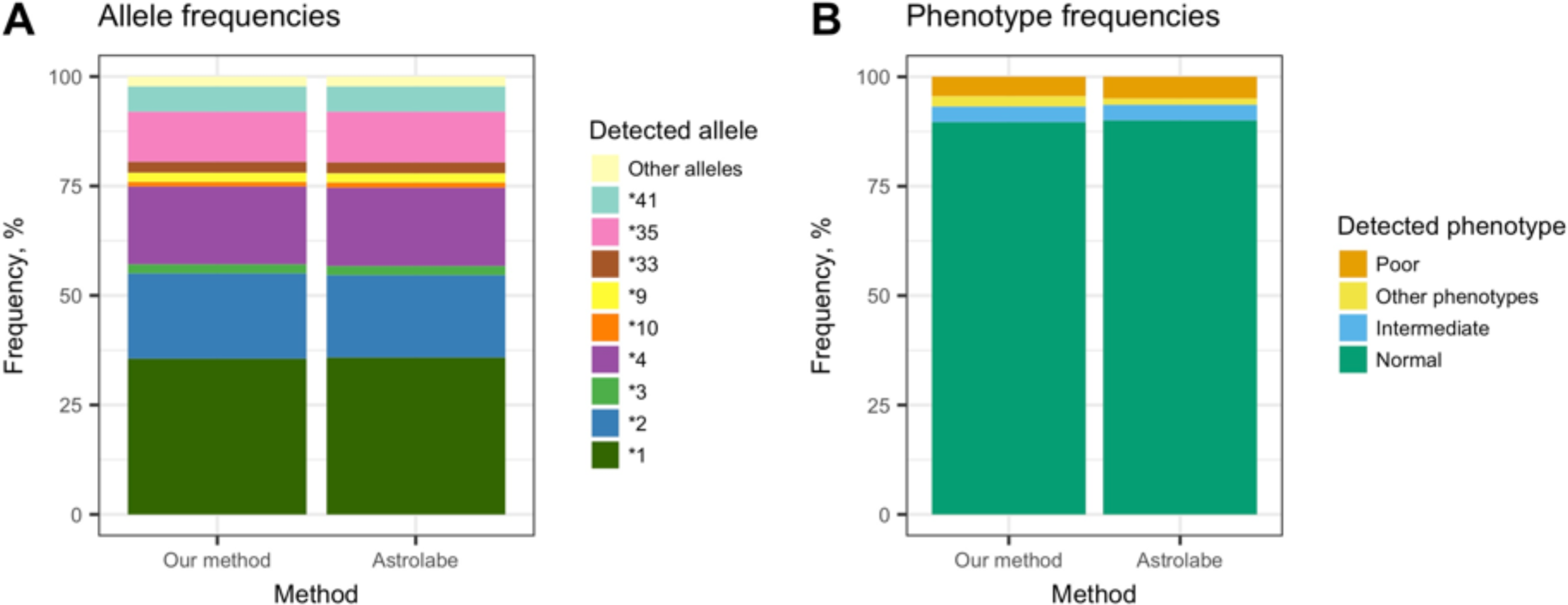
CYP2D6 allele and phenotype frequencies in WGS derived by two methods (our method, Astrolabe)

## REFERENCES

1. Lauschke, V. M., Milani, L. & Ingelman-Sundberg, M. Pharmacogenomic Biomarkers for Improved Drug Therapy—Recent Progress and Future Developments. AAPS J. 20, 4 (2018).

2. Lauschke, V. M. & Ingelman-Sundberg, M. Requirements for comprehensive pharmacogenetic genotyping platforms. Pharmacogenomics 17, 917–924 (2016).

3. Kozyra, M., Ingelman-Sundberg, M. & Lauschke, V. M. Rare genetic variants in cellular transporters, metabolic enzymes and nuclear receptors can be important determinants of interindividual differences in drug response. Genet. Med. 19, 20–29 (2017).

4. Rasmussen-Torvik, L. J. et al. Design and Anticipated Outcomes of the eMERGE-PGx Project: A Multicenter Pilot for Preemptive Pharmacogenomics in Electronic Health Record Systems. Clin. Pharmacol. Ther. 96, 482–489 (2014).

5. Dunnenberger, H. M. et al. Preemptive clinical pharmacogenetics implementation: current programs in five US medical centers. Annu. Rev. Pharmacol. Toxicol. 55, 89–106 (2015).

6. van der Wouden, C. et al. Implementing Pharmacogenomics in Europe: Design and Implementation Strategy of the Ubiquitous Pharmacogenomics Consortium. Clin. Pharmacol. Ther. 101, 341–358 (2017).

7. Caudle, K. E. et al. Incorporation of pharmacogenomics into routine clinical practice: the Clinical Pharmacogenetics Implementation Consortium (CPIC) guideline development process. Curr. Drug Metab. 15, 209–17 (2014).

8. Robarge, J. D., Li, L., Desta, Z., Nguyen, A. & Flockhart, D. A. The Star-Allele Nomenclature: Retooling for Translational Genomics. Clin. Pharmacol. Ther. 82, 244–248 (2007).

9. Kalman, L. et al. Pharmacogenetic allele nomenclature: International workgroup recommendations for test result reporting. Clin. Pharmacol. Ther. 99, 172–185 (2016).

10. Klein, T. E. & Ritchie, M. D. PharmCAT: A Pharmacogenomics Clinical Annotation Tool. Clin. Pharmacol. Ther. (2017). doi:10.1002/cpt.928

11. Leitsalu, L., Alavere, H., Tammesoo, M.-L., Leego, E. & Metspalu, A. Linking a Population Biobank with National Health Registries—The Estonian Experience. J. Pers. Med. 5, 96–106 (2015).

12. Li, H. & Durbin, R. Fast and accurate short read alignment with Burrows-Wheeler transform. Bioinformatics 25, 1754–1760 (2009).

13. McKenna, A. et al. The Genome Analysis Toolkit: a MapReduce framework for analyzing next-generation DNA sequencing data. Genome Res. 20, 1297–303 (2010).

14. Van der Auwera, G. A. et al. in Current Protocols in Bioinformatics 43, 11.10.1-11.10.33 (John Wiley & Sons, Inc., 2013).

15. Danecek, P. et al. The variant call format and VCFtools. Bioinformatics 27, 2156–8 (2011).

16. Mitt, M. et al. Improved imputation accuracy of rare and low-frequency variants using population-specific high-coverage WGS-based imputation reference panel. Eur. J. Hum. Genet. 25, 869–876 (2017).

17. Li, H. & Wren, J. Toward better understanding of artifacts in variant calling from high-coverage samples. Bioinformatics 30, 2843–2851 (2014).

18. Purcell, S. et al. PLINK: A Tool Set for Whole-Genome Association and Population-Based Linkage Analyses. Am. J. Hum. Genet. 81, 559–575 (2007).

19. Loh, P.-R., Palamara, P. F. & Price, A. L. Fast and accurate long-range phasing in a UK Biobank cohort. Nat. Genet. 48, 811–816 (2016).

20. Loh, P.-R. et al. Reference-based phasing using the Haplotype Reference Consortium panel. Nat. Genet. 1443–1448 (2016).

21. Browning, B. L. & Browning, S. R. A Unified Approach to Genotype Imputation and Haplotype-Phase Inference for Large Data Sets of Trios and Unrelated Individuals. Am. J. Hum. Genet. 84, 210–223 (2009).

22. Jia, X. et al. Imputing Amino Acid Polymorphisms in Human Leukocyte Antigens. 8, (2013).

23. Handsaker, R. E. et al. Large multiallelic copy number variations in humans. Nat. Genet. 47, 296–303 (2015).

24. Gaedigk, A., Sangkuhl, K., Whirl-Carrillo, M., Klein, T. & Leeder, J. S. Prediction of CYP2D6 phenotype from genotype across world populations. Genet. Med. 19, 69–76 (2017).

25. R Core Team. R Core Team (2017). R: A language and environment for statistical computing. R Found. Stat. Comput. Vienna, Austria. URL http://www.R-project.org/. R Foundation for Statistical Computing (2017).

26. Wickham, H. & Francois, R. dplyr: A Grammar of Data Manipulation. R Packag. version 0.4.2. 3 (2015). doi:10.18637/jss.v072.i07>.Depends

27. Wickham, H. Reshaping Data with the reshape Package. J. Stat. Softw. 21, 1–20 (2007).

28. Wickham, H. Ggplot2. Elegant Graphics for Data Analysis (2009). doi:10.1007/978-0-387-98141-3

29. Twist, G. P. et al. Constellation: a tool for rapid, automated phenotype assignment of a highly polymorphic pharmacogene, CYP2D6, from whole-genome sequences. NPJ genomic Med. 1, 15007 (2016).

30. Phillips, E. J. et al. Clinical Pharmacogenetics Implementation Consortium Guideline for HLA Genotype and Use of Carbamazepine and Oxcarbazepine: 2017 Update. Clin. Pharmacol. Ther. 103, 574–581 (2018).

31. Saito, Y. et al. CPIC: Clinical Pharmacogenetics Implementation Consortium of the Pharmacogenomics Research Network. Clin. Pharmacol. Ther. 99, 36–37 (2016).

32. Martin, M. A. et al. Clinical pharmacogenetics implementation consortium guidelines for HLA-B genotype and abacavir dosing: 2014 update. Clin. Pharmacol. Ther. 95, 499–500 (2014).

33. Muir, A. J. et al. Clinical Pharmacogenetics Implementation Consortium (CPIC) Guidelines for IFNL3 (IL28B) Genotype and PEG Interferon-α-Based Regimens. Clin. Pharmacol. Ther. 95, 141–146 (2014).

34. Fricke-Galindo, I. et al. Interethnic variation of CYP2C19 alleles, ‘predicted’ phenotypes and ‘measured’ metabolic phenotypes across world populations. Pharmacogenomics J. 16, 113–123 (2016).

35. Gaedigk, A. et al. The CYP2D6 Activity Score: Translating Genotype Information into a Qualitative Measure of Phenotype. Clin. Pharmacol. Ther. 83, 234–242 (2008).

36. Samwald, M., Blagec, K., Hofer, S. & Freimuth, R. R. Analyzing the potential for incorrect haplotype calls with different pharmacogenomic assays in different populations: a simulation based on 1000 Genomes data. Pharmacogenomics 16, 1713–1721 (2015).

37. Nelson, M. R. et al. An Abundance of Rare Functional Variants in 202 Drug Target Genes Sequenced in 14,002 People. Science (80-.). 337, 100–104 (2012).

38. Gordon, A. S. et al. Quantifying rare, deleterious variation in 12 human cytochrome P450 drug-metabolism genes in a large-scale exome dataset. Hum. Mol. Genet. 23, 1957–1963 (2014).

39. Fujikura, K., Ingelman-Sundberg, M. & Lauschke, V. M. Genetic variation in the human cytochrome P450 supergene family. Pharmacogenet. Genomics 25, 584–94 (2015).

40. Bush, W. et al. Genetic variation among 82 pharmacogenes: The PGRNseq data from the eMERGE network. Clin. Pharmacol. Ther. 100, 160–169 (2016).

41. Han, S. et al. Targeted Next-Generation Sequencing for Comprehensive Genetic Profiling of Pharmacogenes. Clin. Pharmacol. Ther. 101, 396–405 (2017).

42. Ahn, E. & Park, T. Analysis of population-specific pharmacogenomic variants using next-generation sequencing data. Sci. Rep. 7, 8416 (2017).

43. Mizzi, C. et al. Personalized pharmacogenomics profiling using whole-genome sequencing. Pharmacogenomics 15, 1223–1234 (2014).

44. Bush, W. S. et al. Genetic variation among 82 pharmacogenes: The PGRNseq data from the eMERGE network. Clin. Pharmacol. Ther. 160–169 (2016). doi:10.1002/cpt.350

45. Lauschke, V. M. & Ingelman-Sundberg, M. Precision Medicine and Rare Genetic Variants. Trends Pharmacol. Sci. 37, 85–6 (2015).

46. Lauschke, V. M. & Ingelman-Sundberg, M. How to Consider Rare Genetic Variants in Personalized Drug Therapy. Clin. Pharmacol. Ther. 103, 745–748 (2018).

47. Yang, W. et al. Comparison of genome sequencing and clinical genotyping for pharmacogenes. Clin. Pharmacol. Ther. 100, 380–388 (2016).

48. McKinnon, R. A., Ward, M. B. & Sorich, M. J. A critical analysis of barriers to the clinical implementation of pharmacogenomics. Ther. Clin. Risk Manag. 3, 751–9 (2007).

49. Maciel, A., Cullors, A., Lukowiak, A. A. & Garces, J. Estimating cost savings of pharmacogenetic testing for depression in real-world clinical settings. Neuropsychiatr. Dis. Treat. 14, 225–230 (2018).

50. Winner, J., Allen, J. D., Anthony Altar, C. & Spahic-Mihajlovic, A. Psychiatric pharmacogenomics predicts health resource utilization of outpatients with anxiety and depression. Transl. Psychiatry 3, e242–e242 (2013).

51. Chou, W. H. et al. Extension of a pilot study: impact from the cytochrome P450 2D6 polymorphism on outcome and costs associated with severe mental illness. J. Clin. Psychopharmacol. 20, 246–51 (2000).

52. Van Driest, S. L. et al. Clinically actionable genotypes among 10,000 patients with preemptive pharmacogenomic testing. Clin. Pharmacol. Ther. 95, 423–431 (2014).

53. Caraballo, P. J. et al. Multidisciplinary model to implement pharmacogenomics at the point of care. Genet. Med. 19, 421–429 (2017).

